# Behavioural variation in background colour matching and the effect of social environment in rock-pool mosquitoes

**DOI:** 10.1101/2024.07.24.604960

**Authors:** Giulia Cordeschi, Daniele Porretta, Daniele Canestrelli

**Affiliations:** Department of Environmental Biology, Sapienza University of Rome, Italy; Department of Ecology and Biology, Tuscia University, Viterbo, Italy

## Abstract

Several animal species conceal themselves from potential predators by actively choosing environmental patches that best match body colouration and chromatic patterns. Growing evidence shows that a variety of contexts and conditions can affect this background choice behaviour, yet the extent of this variety remains largely underexplored. Here, we explore the effect of disturbance and social environment on background colour choice behaviour in the rock-pool mosquito *Aedes mariae*. We exposed single individuals and groups of individuals to experimental arenas made of dark and bright patches, and recorded individuals’ colour preferences when alone and within groups, as well as before and after a disruptive event. We found a marked prevalence of individuals favouring a dark background and an among-individual variation in choice over replicated trials. Moreover, we observed a non-significant effect of disturbance but a significant role of the social environment. In fact, being caged in groups significantly increased the proportion of mosquitoes choosing the dark background. Our results provide strong evidence of a background colour choice in *Ae. mariae*, a density-dependence of this choice, and a non-negligible inter-individual variation in this behaviour. Overall, these findings offer intriguing insights into the background choice behavioural variation and flexibility in mosquitoes.

## Introduction

The ability of many animal species to conceal themselves from predators by resembling the surrounding environment has long attracted scientific curiosity (Ruxton et al. 2004, Merilaita and Lind 2005, Merilaita and Stevens 2011, Stevens and Ruxton 2019, Pembury Smith and Ruxton 2020). An effective crypsis can be achieved with two main strategies. Masters of camouflage, such as chameleons and cephalopods, can change their body colour in seconds, enabling them to fine-tune their resemblance to a background that changes while the animal moves (Hanlon 2007, Pembury Smith and Ruxton 2020). However, many animals do not possess physiologically mediated colour change abilities, and behaviour can be critical in achieving effective crypsis in these cases (Stevens and Ruxton 2019). The most obvious way that animals could use behaviour to improve camouflage is by choosing to rest on the backgrounds that best match their appearance, i.e. background choice (Merilaita and Stevens 2011). Examples of background choice behaviour date back to the observation of A.R. Wallace, who noticed that the effectiveness of the camouflage of the Indian leaf butterfly, *Kallima inachus*, resides on the choice of resting on dead leaves and twigs, but not on fresh green vegetation (Wallace 1867). In his pioneering study on background matching in the peppered moth *Biston betularia*, Kettlewell (1955) used alternate black and white paper stripes to investigate background choice behaviour of the two different morphs, the pale and the melanic form (Kettlewell 1955, Eacock et al. 2019). He observed marked preferences for a dark background by the melanic, while the opposite preference was observed for the pale morph (Kettlewell 1955).

Density-dependent processes can regulate background matching components (Tollrian et al. 2015). The defence level often increases with predator density, and it was observed that predation risk increases the likelihood of behavioural background matching (Edelaar et al. 2017). In Aegean wall lizards (*Podarcis erhardii*), individuals chose a background that resembled their dorsal colouration, and this pattern is more marked on islands with higher predation risk (Marshall et al. 2016). By analyzing background choice before and after a disturbance in tadpoles of the treefrog *Bokermannohyla alvarengai*, Eterovick and colleagues (2018) observed that tadpoles move on a substrate that resembles their brightness and colour, improving camouflage when disturbed. On the other hand, individual predation risk depends not only on predator density but also on the density of conspecifics. Although one of the better-known effects of aggregation is to reduce individual predation risk (Krause and Ruxton 2002), detection risk generally increases with group size (Krause and Godin 1995, Jackson et al. 2005, Ioannou and Krause 2008). Several studies on raptors attacking flocks of birds have found a positive correlation between the number of attacks and the size of the prey group (Lindstrom 1989, Cresswell 1994). Studying the effect of group size on detection rate in a field experiment, Ioannou and Krause (2008) found that three-spined sticklebacks *Gasterosteus aculeatus* take longer time to approach and attack a group of two *Daphnia*, compared to a group of a hundred individuals (Ioannou and Krause 2008). However, while the effect of the social environment on per-capita predation risk has been largely investigated, we are still virtually blind to its effects on background choice behaviours.

In this paper, we investigated background choice behaviour and the effect of disturbance and conspecific density in the sea rock-pool mosquito *Aedes mariae*. This species is distributed along the western Mediterranean coasts, developing in rock-pools of the supralittoral zone of coastal habitats. Adult mosquitoes have the ability to discriminate between colours, with some differences between species, and colour preferences were observed in the behaviour of host-seeking, oviposition and feeding (Muir et al. 1992, Bidlingmayer 1994, Briscoe and Chittka 2001, S A Allan et al. 2003, Jung et al. 2021), but not in the highly predation-relevant resting behaviour. Here, we address (i) the background colour preference in resting behaviour in rock-pool mosquitoes, (ii) the effect of a potential threat, and (iii) the effect of the social environment on background choice behaviour. We hypothesized that mosquitoes might prefer to rest on a dark background to best match the surrounding colour and to decrease the probability of being detected, and that this behaviour might be flexible in response to disturbance and conspecific density.

## Materials and Methods

We carried out experiments on a total of 200 adults of *Ae. mariae* obtained from larvae collected in July 2022 from supralittoral rock-pools of San Felice Circeo, central-western Italy (41°13’18.77”N, 13°04’05.51”E). Larvae were reared in plastic trays (5*30*19 cm) filled with water from the sampling pools and fed ad libitum with catfood Friskies®Adult. After adult emergence, mosquitoes were placed individually in plastic containers and fed with 10% sucrose solution. We carried out four biological replicates, testing 50 individuals per replicate.

The experimental arena designed to test background choice behaviour was a cubic cage (45*45*45 cm) with three side walls and the roof covered with a chessboard of dark and light patches. The front wall was made of transparent plastic to allow behavioural observations.

To assess the individual colour preference, mosquitoes were placed individually inside the experimental arena with the home cage/container. The individual was let out of the home cage spontaneously after 10 minutes of acclimation, and the choice of background colour to rest on was recorded (Fig. 1). Each individual was tested twice on two consecutive days.

**Figure 1:**
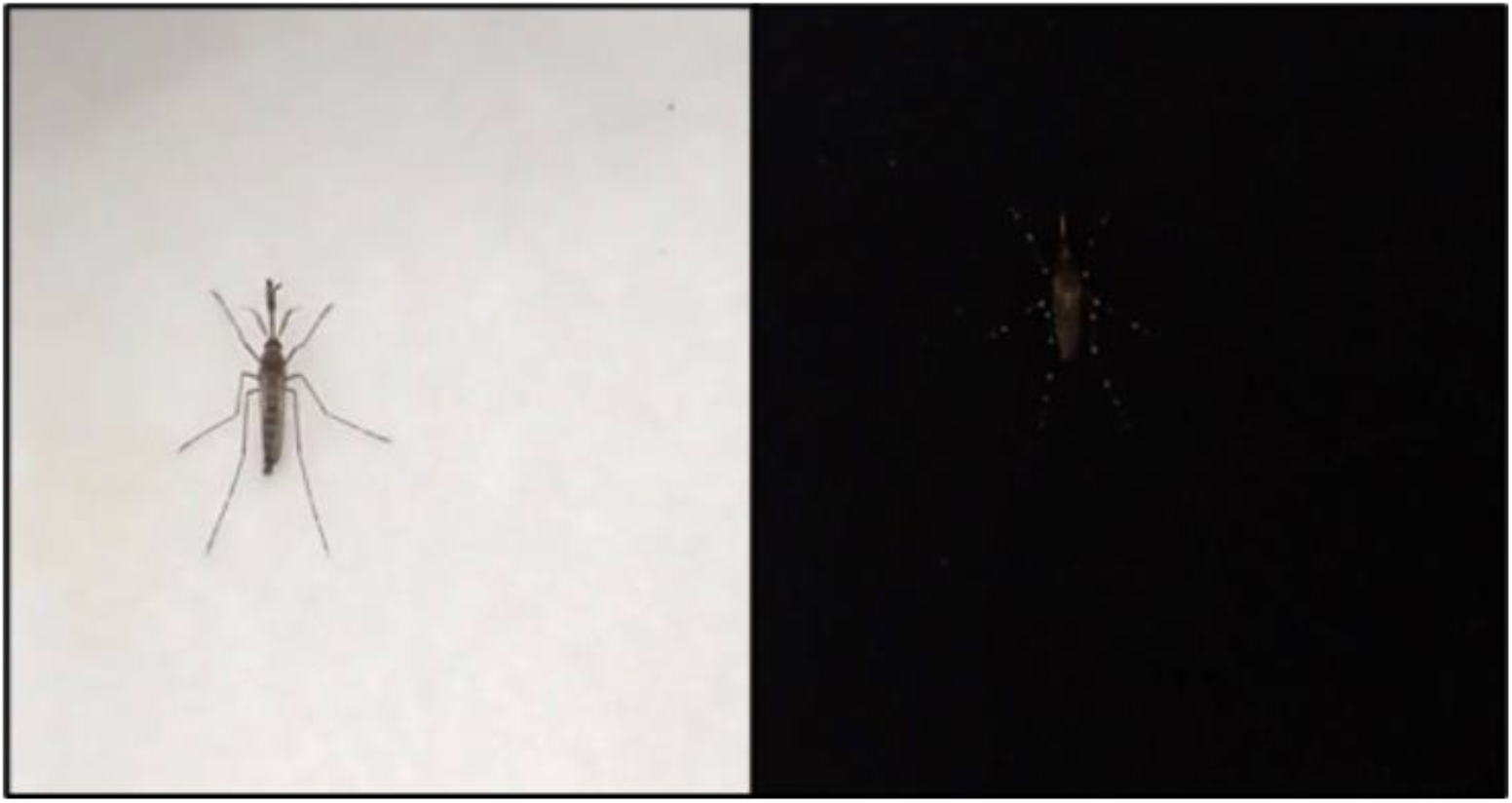
Appearance of a male rock-pool mosquito *Aedes mariae* on light and dark backgrounds

Because predation risk can increase the likelihood of behavioural background matching (see e.g. Eterovick et al. 2018), we investigated the effect of a potential threat on this behaviour. When the individual stopped on the artificial background, a disturbance was provided by gently tapping on the walls of the experimental cage, and the subsequent colour choice was recorded. Each individual was tested twice on two consecutive days.

To evaluate the effect of social environment, we tested the same individuals in groups of 50 mosquitoes, recording the number of individuals on each colour twice without the disturbance stimulus and twice after the disturbance.

### Statistical analysis

Generalized linear mixed-effect models (GLMMs) were used to analyze individual-level variation in background choice behaviour. The analysis was run using the *glmer* function in the *lme4* package (Bates et al. 2015) with a Binomial distribution and logit link function. We included disturbance as a fixed effect, while individual identity was included as a random effect (random intercept). We assessed the importance of the contribution of consistent individual differences to total phenotypic variance by determining the significance of the random effect of individual mosquito ID using likelihood ratio tests (LRTs) and by comparing Akaike’s Information Criterion (AICs) for models with and without the random effect. Likelihood ratio tests were also performed to calculate the significance of fixed effects included in the model. To check for differences between biological replicates, we included it as a fixed effect in the initial model, but it was not found to significantly affect background choice behaviour (*χ*^*2*^ = 5.68, *p* = 0.459) and was removed from the final model.

To test whether social environment influences background choice behaviour, we applied GLM with binomial distribution using the number of individuals choosing dark and the number of individuals choosing light backgrounds as dependent variables and conspecific density (individual vs group), condition (with vs without disturbance stimulus) ad their interaction as fixed effects. We performed Likelihood ratio test to calculate the significance of fixed effects and pair-wise comparisons using *emmeans* package with the conservative Bonferroni method for significant predictors, accounting for the variation explained by other predictors (Lenth 2022). To check for the effect of id of groups, we included it as a fixed effect in the initial model, but it was not found to significantly affect background choice behaviour (*χ*^*2*^ = 1.658, *p* = 0.197) and was removed from the final model. We analyzed the data in R version 4.1.2 (R Core Team, 2016).

## Results

Results showed an overall marked preference for dark backgrounds in both individual and group tests and with or without disturbance (Fig. 2). In individual tests we found a substantial among-individual variation (Tab. 1) while disturbance stimulus did not significantly influence the background choice behaviour (Tab. 1). We observed that without the disturbance stimulus, in the first trial 72.5% and in the second trial 80% of mosquitoes chose dark backgrounds while after the disturbance 74 % in the first trial and 82% in the second trial. This pointed out that there wasn’t a directional change towards the choice of a dark background following a potentially threatening stimulus.

**Tab.1:** Results from GLMMs

| **Model: Individual choice test** |  |  |  |  |
| --- | --- | --- | --- | --- |
| **Fixed effect** | **Estimate ± SE** |  | **X<sup>2</sup>** | ***p*** |
| Disturbance | 0.111 ± 0.178 | | 0.388 | 0.533 |
| **Random effect (ID)** | **Estimate ± SE** | **ΔAIC** | **X<sup>2</sup>** | ***p*** |
| $V_{\text{among}}$ | 0.699 ± 0.836 | 10.78 | 12.776 | **<0.001** |
| $V_{\text{within}}$ | 5.688 ± 0.313 | | | |
| **Model: Group choice test** |  |  |  |  |
| **Fixed effect** | **Estimate ± SE** |  | **X<sup>2</sup>** | ***p*** |
| Disturbance | 0.454 ± 0.221 | | 4.6393 | 0.098 |
| Density | 0.899 ± 0.207 | | 28.921 | **<0.001** |
| Disturbance*Density | 0.352 ± 0.278 | | 1.640 | 0.200 |

**Figure 2:**
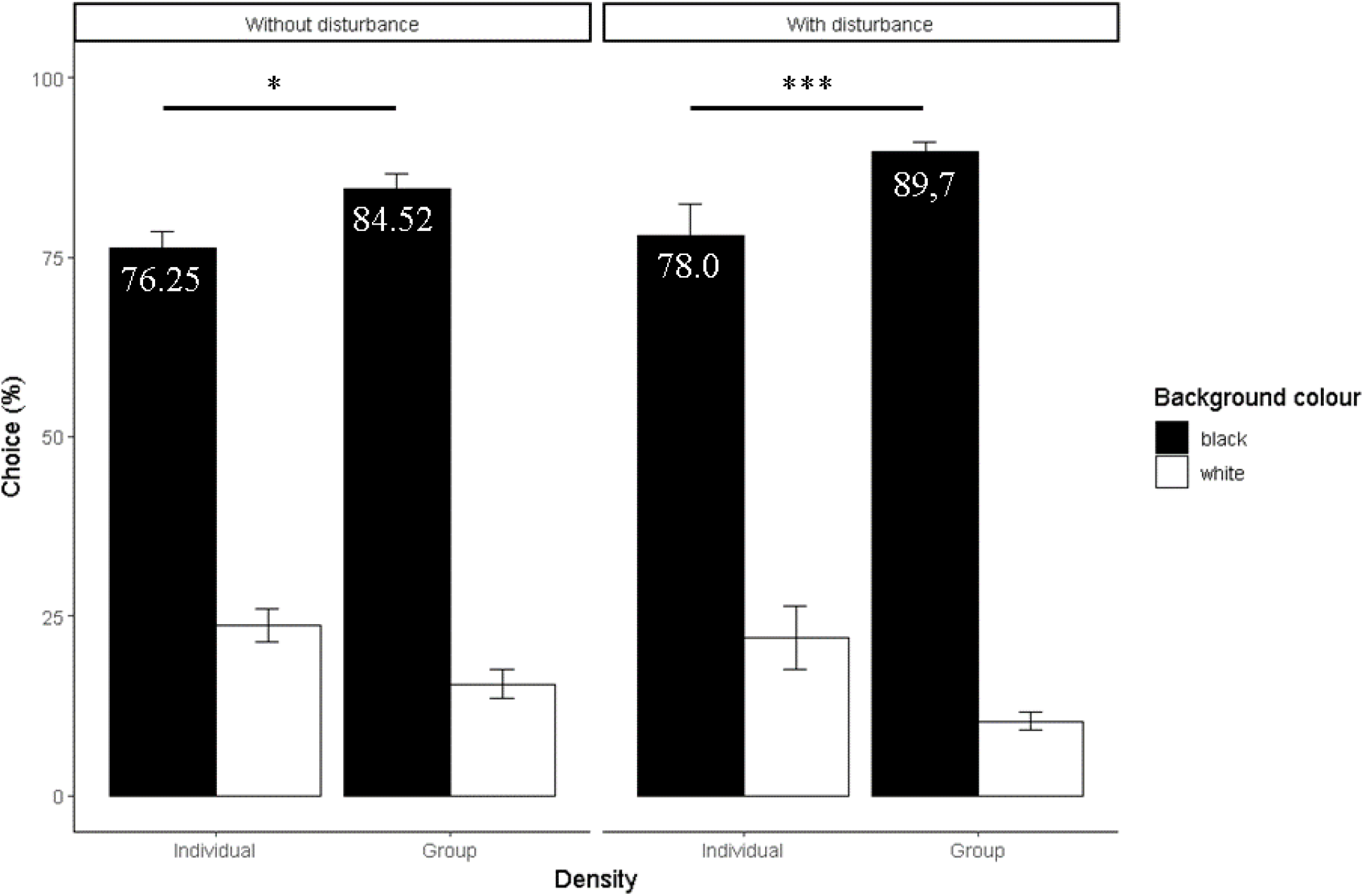
Background colour choice before (left) and after a disturbance stimulus (right), in individual and group tests. Mean between repeated measures ±s.e. Significance levels of multiple comparisons: *** = P < 0.001, * = P < 0.05.

On the other hand, social environment had a significant overall effect on background choice behaviour (figure 2; Tab. 1). Comparing individual versus group choices, we found a significant increment of individuals choosing dark backgrounds both without disturbance (Individual tests: 76.25± 2.28 %; Group tests: 84.52 %, trials mean ± se) and after disturbance (Individual tests: 78.0 ± 4.41%%; Group tests: 89.7± 1.27 %, trials mean ± se) (Fig. 2). As for individual tests, disturbance had no significant effect neither the interaction between conspecific density and disturbance (Tab. 1).

## Discussion

Vision plays an essential role in several behaviours of mosquito species, and their colour preference mediates host-seeking behaviour, oviposition site selection and foraging (Allan et al. 1987, Takken and Verhulst 2013, Jung et al. 2021, Alonso San Alberto et al. 2022, Hellhammer et al. 2022). In *Aedes albopictus* and *Culex pipiens*, dark colours such as black, red, blue and purple are more attractive than light colours during host-seeking behaviour (Browne and Bennett 1981, Jung et al. 2021, Alonso San Alberto et al. 2022). In *Anopheles, Aedes* and *Culex* female mosquitoes, a preference for black, red, or blue was also observed in the context of oviposition site selection (Hoel et al. 2011, Hellhammer et al. 2022, Phiri and Mbata 2023, Moazzeni et al. 2023). Most studies of mosquito colour preferences have been carried out using other attractants as host olfactory stimuli (but see Hellhammer et al. 2022) and were conducted primarily on vector species to develop effective control strategies or advanced traps (Bidlingmayer 1994, Hoel et al. 2011). Here, we explored background choice behaviour as an antipredator strategy in adult mosquitoes and the effect of social environment and disturbance, without using other attractants. We found a strong preference for dark backgrounds in resting behaviour in *Aedes mariae*. Thermoregulation could be linked to background colour choice (Shreeve 1990), but since we used the same material for both colour patches and conducted the experiments in a climatic chamber, it seems unlikely that temperature conditions drove the choice. On the other hand, resting on a dark surface, mosquitoes reduced substantially the chromatic contrast between their body and the surroundings (Fig.1), thus limiting the probability of being detected by a predator.

Anti-predator behaviour can vary between contexts, although the extent of this variation has been poorly characterized, particularly for background matching behaviours. We found that the choice of *Ae. mariae* adults for dark or pale backgrounds was not affected by a potentially threatening stimulus, while the choice of dark surfaces was affected by the social context. Indeed, when housing mosquitoes in large groups, the number of individuals choosing the dark background was markedly higher than when they were housed individually. Although we did not assess individual personality and their potential association with background colour choice, this higher preference for a darker background might in fact suggest an attitude toward a more prudent behaviour when in groups. Group size has been found to modulate antipredator behaviour in a wide range of insects, usually with bolder behaviours observed when individuals are in a group (Brown et al. 2006, Smith and Awan 2009). When exposed to a predatory stimulus, individuals of the marine insect *Halobates robustus*, alternatively show a ‘confusion’ or an escape behaviour when housed in large or small groups, respectively (Treherne and Foster 1982). In *Ae. mariae*, the increased background matching behaviour triggered by a high density of conspecifics might be associated with a higher risk of being detected and predated when aggregated. *Ae. mariae*, as many other culicine species, usually aggregate in the context of large mating swarms (Clements 1999; Takken et al. 2006). Our results that large (non-reproductive) aggregates promote marked preferences for darker backgrounds suggests the hypothesis that during the mating swarms, there may be perceived trade-offs between reproductive and antipredator behaviours (Sullivan 1981, Yuval and Bouskila 1993, Jackson et al. 2005, Ioannou and Krause 2008).

Despite the obvious importance of the inter-individual level of variation for behavioural evolution (Dingemanse and Réale 2005, Bell et al. 2009, Wolf and Weissing 2012), most studies on background matching have concentrated on responses at the between-species or between-morph level of variation, with very limited exploration into how individuals’ choice behaviour vary within populations (Stevens and Ruxton 2019). Here, we observed a high inter-individual variation in background choice behaviour despite the absence of body colour polymorphisms. This variability could be attributed to conflicting evolutionary pressures and ecological trade-offs. Behavioural responses to predation risk are critical for survival but involve trade-offs in terms of energy and time allocation to other essential behaviours like feeding and mating (Sih 1997, Martin and Lopez 1999, Martín et al. 2003b, Reaney 2007). Individuals could modulate their defensive response by taking into account costs and benefits (Martín et al. 2003a, Amo et al. 2007, Citadini and Navas 2013, Ibáñez et al. 2014). Future research should delve deeper into how anti-predatory behaviour might be shaped by individual personality, individual condition and the effect of the developmental environment within mosquito populations.

